# HSF2 Co-regulates Protein-coding and Long Non-coding RNA Genes Specific to Black Tissues of the Black Chicken, Yeonsan Ogye

**DOI:** 10.1101/223644

**Authors:** Hyosun Hong, Han-Ha Chai, Kyoungwoo Nam, Dajeong Lim, Kyung-Tai Lee, Yoon Jung Do, Chang-Yeon Cho, Jin-Wu Nam

**Affiliations:** Department of Life Science, College of Natural Sciences, Hanyang University, Seoul 133791, Republic of Korea; Research Institute for Convergence of Basic Sciences, Hanyang University, Seoul 133791, Republic of Korea; Department of Animal Biotechnology & Environment of National Institute of Animal Science, RDA, Wanju 55365, Republic of Korea; College of Pharmacy, Chonnam National University, Kwangju 61186, Republic of Korea; Animal Genetic Resource Research Center of National Institute of Animal Science, RDA, Namwon 55717, Republic of Korea

## Abstract

The Yeonsan Ogye (Ogye) is a rare Korean domestic chicken breed, the entire body of which, including its feathers and skin, has a unique black coloring. Although some protein-coding genes related to this unique feature have been examined, non-coding elements have not been globally investigated. In this study, high-throughput RNA sequencing and DNA methylation sequencing were performed to dissect the expression landscape of 14,264 Ogye protein-coding and 6900 long non-coding RNA (lncRNA) genes along with DNA methylation landscape in twenty different Ogye tissues. About 75% of Ogye lncRNAs showed tissue-specific expression whereas about 45% of protein-coding genes did. For some genes, the tissue-specific expression levels were inversely correlated with DNA methylation levels in their promoters. About 39% of the tissue-specific lncRNAs displayed functional association with proximal or distal protein-coding genes. In particular, heat shock transcription factor 2 (HSF2)-associated lncRNAs were discovered to be functionally linked to protein-coding genes that are specifically expressed in black skin tissues, tended to be more syntenically conserved in mammals, and were differentially expressed in black tissues relative to white tissues. Our results not only facilitate understanding how the non-coding genome regulates unique phenotypes but also should be of use for future genomic breeding of chickens.

## Introduction

The Yeonsan Ogye (Ogye) chicken is one of the rarest breeds of *Gallus gallus domesticus*. Domesticated in the Korean peninsula, it probably originated from the Indonesian Ayam Cemani black chicken, which populates tropical, high-temperature areas ^1^. Ogye shares common features—such as black plumage, skin, shank, and fascia—with Ayam Cemani ^1^, although it has a smaller comb and shorter legs. Silkie fowl (Silkie), one of the most popular black-bone chickens, also has black skin but has white or varied color plumage ^2^. Several genes involved in Silkie skin hyperpigmentation have been reported in previous studies ^2-4^. Recently, transcriptomes from Chinese native black chickens were compared with those from white chickens to globally identify hyperpigmentation-related genes ^5^. However, studies of the molecular mechanisms and pathways related to black chicken hyperpigmentation have been restricted to coding genes.

A major part of the non-coding transcriptome corresponds to long non-coding RNAs (lncRNAs), which originate from intergenic, intervening, or antisense-overlapping regions of protein-coding genes ^6-8^. lncRNAs are defined as transcripts longer than 200nt and are mostly untranslated because they lack an open reading frame; however, they interact with RNA binding proteins and have diverse intrinsic RNA functions ^9-11^. They tend to be localized to subcellular areas, particularly the nucleus, and often interact with heterochromatin remodelers and DNA methylation regulators to regulate gene expression at the epigenetic level. For instance, DNMT1-associated colon cancer repressed lncRNA-1 (DACOR1) is localized to genomic sites, known to be differentially methylated, and regulates methylation at least 50 CpG sites by recruiting DNMT1 in colon cancers ^12^.

lncRNAs are also known to regulate gene expression at other levels: transcriptional, post-transcriptional, translational, and post-translational ^9,10,13-15^. They regulate distant genes by modulating the recruitment of transcription factors (TFs) to target genes. Only a few lncRNAs, however, have been experimentally validated as functional; most candidates remain unvalidated. In particular, some lncRNAs have been shown to regulate the expression of neighboring genes in a *cis*-acting manner ^16-20^. Enhancer-associated lncRNAs (eRNAs) are a well-known group in this class that regulate the expression of downstream genes. Knockdown of eRNAs reduces target gene expression, suggesting their function as *cis*-acting elements ^21-23^. eRNA regulatory roles are known to be achieved via several mechanisms: trapping transcription factors, directing chromatin roofing, and inducing DNA methylation ^9,24-28^. On the other hand, lncRNAs that associate with post-transcriptional regulators control target splicing and stability. For instance, antisense lncRNA from the *FGFR2* locus promotes cell-type specific alternative splicing of FGFR2 by interacting with polycomb complex ^29^.

Despite their regulatory roles, only a few lncRNAs are highly conserved across vertebrates ^30^. lncRNAs generally exhibit either poor conservation at the nucleotide level or conservation in a short region only, particularly compared to protein-coding genes ^31-33^. Although sequence conservation is often likely to indicate related function, sometimes it is difficult to detect conservation across multiple genome sequences because of technical challenges. lncRNAs, however, appear to be syntenically conserved with protein-coding genes, which suggests that lncRNAs could have evolutionarily conserved roles in similar genomic contexts ^34-36^. A zebrafish lncRNA, linc-*oip5*, which has a short region of sequence conservation with mammalian orthologs in the last exon, also exhibits preserved genomic architecture in its size and arrangement of exons; furthermore, linc-*oip5* loss of function disrupts zebrafish embryonic development, which can be rescued by the mammalian orthologs ^37^. Thus, examining the genomic context and/or short regions of conservation in a lncRNA may be necessary for understanding lncRNA function.

lncRNA expression signatures also provide hints about lncRNA functional roles at the cellular level. Global lncRNA profiling demonstrated that lncRNAs generally exhibit lower expression than protein-coding genes ^31,38,39^ but tend to be uniquely or specifically expressed in distinct tissues, developmental stages, conditions, or disease states ^30-32,38,40-42^. For instance, one lncRNA, SAMMSON, is specifically expressed in melanoma cells during melanogenesis and is known to regulate the process at the epigenetic level ^43^. In addition, large-scale analyses of lncRNA and protein-coding gene co-expression led to the finding that a considerable number of paired genes are actually co-regulated by common TFs ^44,45^. Often common TF binding motifs have been discovered in the promoters of the co-expressed lncRNA and protein-coding genes, suggesting that the co-regulated genes could share functional roles ^46,47^. Thus, to predict lncRNA biological functions, co-expression networks of lncRNAs and protein-coding genes from large scale transcriptomic data have been constructed and used for the inference of function ^48^-^50^.

Although genome and transcriptome maps of livestock animals, such as rainbow trout, cow, goat, and chicken ^51^-^55^, have been recently constructed, only a few non-coding transcriptome studies have been done in those genomes. To date, 9,681 lncRNAs have been annotated in the red jungle fowl *Gallus gallus* genome, but these studies have been limited to a few tissues. Thus, the expression landscape of Ogye non-coding transcriptomes in many tissues will help us understand phenotypic similarities and differences between Ogye and *Gallus gallus*.

## Results

### Tissue-specific expression and DNA methylation landscapes of Ogye lncRNAs

To profile the expression of lncRNA genes, RNA-seqs were performed across twenty Ogye tissues (Figure 1a; see the “Expression profiling” section in the Methods for more details). Of 6900 Ogye lncRNAs that we annotated in our other study ^56^, 6,565 were expressed with FPKM ≥ 1 in at least one tissue, whereas 13,765 of Ensembl chicken protein-coding genes (release 81; http://www.ensembl.org/biomart) were expressed. Tissue-specific genes with a four-fold higher maximum expression value than the mean value over twenty tissues were depicted on the genome using a Circos plot (Fig. 1b, green track). As previously reported ^32,57,58^ Ogye lncRNAs generally displayed a tissue-specific expression pattern and some lncRNAs were solely expressed in a single tissue, although a few hundreds displayed ubiquitous expression across tissues. About 75% of lncRNA genes (5191 loci) were tissue-specific, a significantly higher proportion than that of protein-coding genes (45%; Fig. 1c; top; Supplementary table S1; *P* < 7.1e^−293^; Fisher’s exact test). The fractions of lncRNAs that were tissue-specific ranged from 2.4 % (Fascia) to 12.5% (Kidney), much higher percentages than those of protein-coding genes, which ranged from 0.4% (Fascia) to 4.2% (Kidney) (Fig. 1d; middle). Hierarchical clustering of commonly expressed lncRNA genes among tissues using the PHYLIP package (ver 3.6) ^59^ (see the “Hierarchical clustering of expressed lncRNAs across tissues” section in the Methods for more details) defined functionally and histologically-related tissue clusters well. In particular, 2,317 lncRNAs were specifically expressed in the comb, skin, and shank, which are black tissues in Ogye (Fig. 1d; left). Only 780 lncRNAs were ubiquitously expressed across all tissues (Fig. 1d; left).

**Figure 1.**
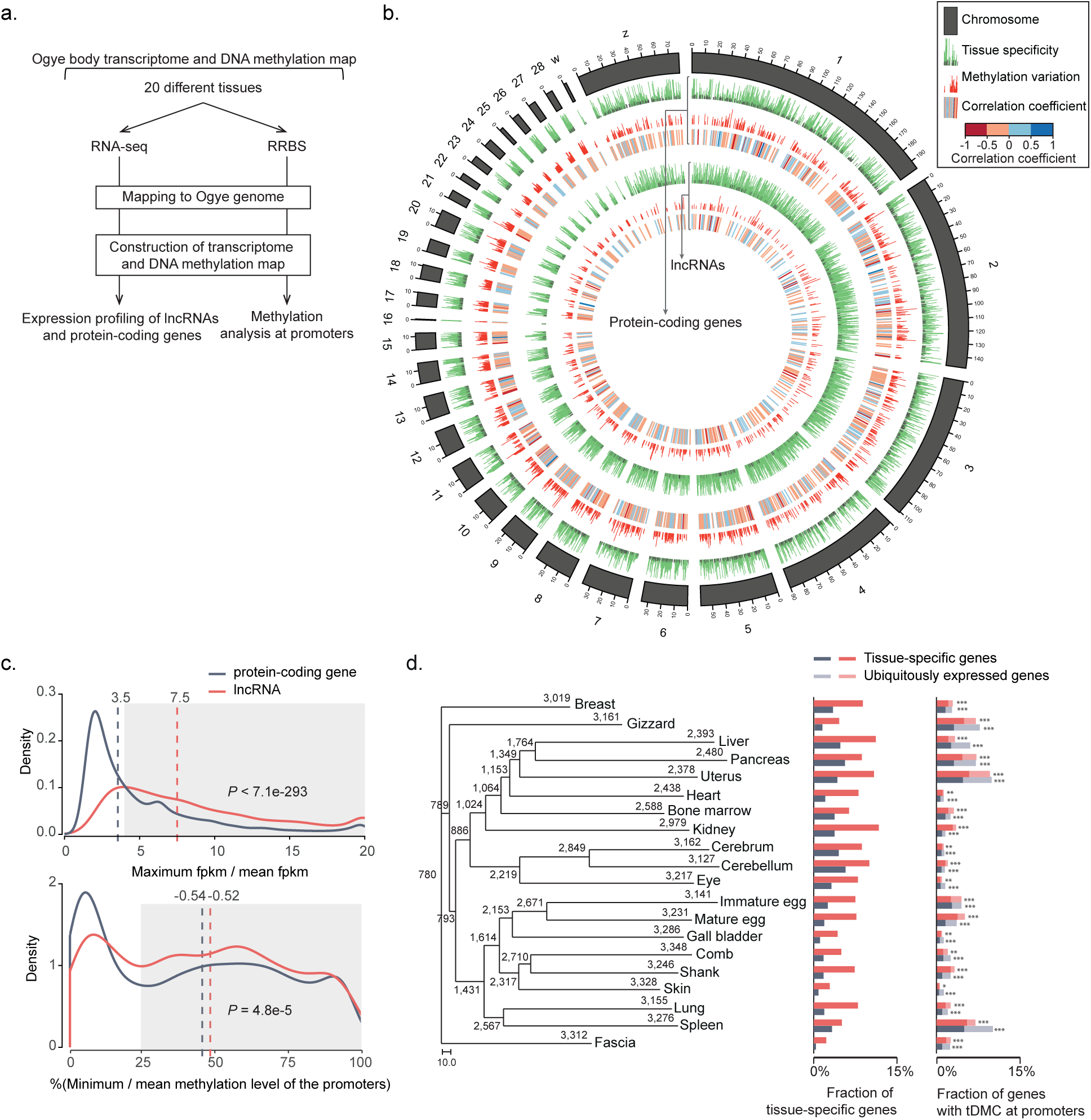
Expression and DNA methylation landscapes of Ogye lncRNAs. **a**. A schematic flow for the analyses of coding and non-coding transcriptomes and DNA methylation from twenty different tissues. **b**. A Circos plot illustrating the expression variability (green bars) of lncRNA and protein-coding genes, the methylation variability (red bars) at tissue-specific, differentially methylated CpG sites in the promoters, and the correlation coefficients between expression and methylation levels across chromosomes (heatmaps). The variability was measured as the ratio of the maximum and mean expression values or methylation levels. The correlation was calculated with Spearman correlation coefficients, which were demonstrated with a heatmap, scaled from −1 to 1. **c**. Shown are the distributions of the maximum versus mean expression values of lncRNA (red line) and protein-coding genes (black line) across tissues (top), and the distributions of the minimum versus mean methylation levels of each cytosine in the promoter of lncRNAs (red line) and protein-coding genes (black line) (bottom). The vertical dotted lines indicate the median value of the respective distribution (black for protein-coding genes and red for lncRNAs). The gray boxes indicate the tissue-specific expression and methylation **d**. Numbers of commonly or uniquely expressed lncRNAs across tissues are shown in the phylogenetic tree of tissues. The numbers at the leaf nodes indicate lncRNAs expressed in the indicated tissue (FPKM ≥ 1) and the numbers at the internal nodes indicate those commonly expressed in the indicated tissues. Of the expressed genes in a certain tissue, the fraction of the tissue-specific genes (red for lncRNA and black for protein-coding genes) and the fraction of genes with a differentially methylated region (DMR) in the promoters are indicated as bar graphs. Of the genes with a DMR, tissue-specific genes (dark) and others (light) were distinguished and the enrichment of the tissue-specific genes was tested using Fisher’s exact test (* *P* ≤ 1 X e^−5^, ** *P* ≤ 1 X e^−10^, *** *P* ≤ 1 X e^−20^). The scale bar represents 10.0, which is the unit of 120 differentially expressed genes across tissues.

To correlate tissue-specific lncRNA expression with its epigenetic status in the respective tissue, the DNA methylation signals were profiled from corresponding tissues using reduced representation bisulfite sequencing (RRBS; see the “Data source” section in the Methods for more details). A significant correlation (nominal *P* ≤ 0.05) between the expression levels and the methylation signals in the region 2kb upstream (regarded as promoter) of genes across twenty tissues was demonstrated along with a mean variation of the signals (Fig. 1b), the ratio of the minimum and mean methylation levels at CpG sites in the corresponding promoter. About 70% of lncRNA genes with CpG methylation signal in their promoter region displayed a tissue-specific methylation with 25% lower minimum methylation value than the mean value over twenty tissues, which is a significantly higher proportion than that of protein-coding genes (64%; Fig. 1c; bottom; *P* = 4.8e^−5^; Fisher’s exact test).

To examine the association between the gene expression and promoter methylation, the genes with tissue-specific differentially methylated CpG sites (tDMC; see the “Tissue-specific, differentially methylated CpG sites” section in the Methods for more details) that include ≥ five reads with C to T changes in ≥ 10 tissues in the promoter region were subjected to the downstream analyses. The fractions of genes that have tDMC in their promoters were significantly enriched in the tissue-specific genes (Fig. 1d; right). Of lncRNA and protein-coding genes with tDMC, 6.4% of the lncRNAs and 9.3% of the protein-coding genes displayed a significant negative correlation (nominal *P* ≤ 0.05) between their promoter methylation levels and their expression levels, percentages that were significantly higher than those of random-pair controls (Supplementary figure S1b; *P* = 1.30 X 10^−6^ for lncRNAs; *P* = 7.93 X 10^−36^ for protein-coding genes; Fisher’s exact test). However, only about 3% of genes showed a positive correlation between their expression and methylation signals, which is comparable or less than the control (Supplementary figure S1c; *P* = 0.87 for lncRNAs; *P* = 0.013 for protein-coding genes). Collectively, these results show that CpG methylation in the promoters is epigenetically associated with the expression of target genes.

### Tissue-specific lncRNA clusters functionally linked to protein-coding genes

As lncRNAs tend to be specifically expressed in a tissue or in related tissues, they could be worthy factors for defining phenotypic characteristics of tissues. To identify functional clusters of lncRNAs, pairwise correlation coefficients between tissue-specific lncRNAs were calculated and the co-expression patterns across twenty tissues were clustered, defining 16 co-expression clusters (Fig. 2). As expected, each co-expression cluster was defined as a functional group, highly expressed in a certain tissue (kidney, eye, pancreas, uterus, mature egg, immature egg, breast, heart, liver, lung, gall bladder, gizzard, bone marrow, or spleen) or related tissues (brain and black tissues) (Supplementary table S2). In particular, the largest co-expression cluster, the brain-specific group, included 930 co-expressed lncRNAs, highly expressed in cerebrum and cerebellum. The second largest cluster, the black tissue-specific group, included 479 co-expressed lncRNAs, highly expressed in fascia, comb, skin, and shank (Fig. 2). Clusters of related tissues also display distinct sub-modules corresponding to each tissue. For instance, lncRNA clusters specific to black tissues displayed sub-clusters including sub-cluster 1 specific to shank and sub-cluster 2 specific to comb, although the sub-clusters shared skin-specific expression (Supplementary figure S2).

**Figure 2.**
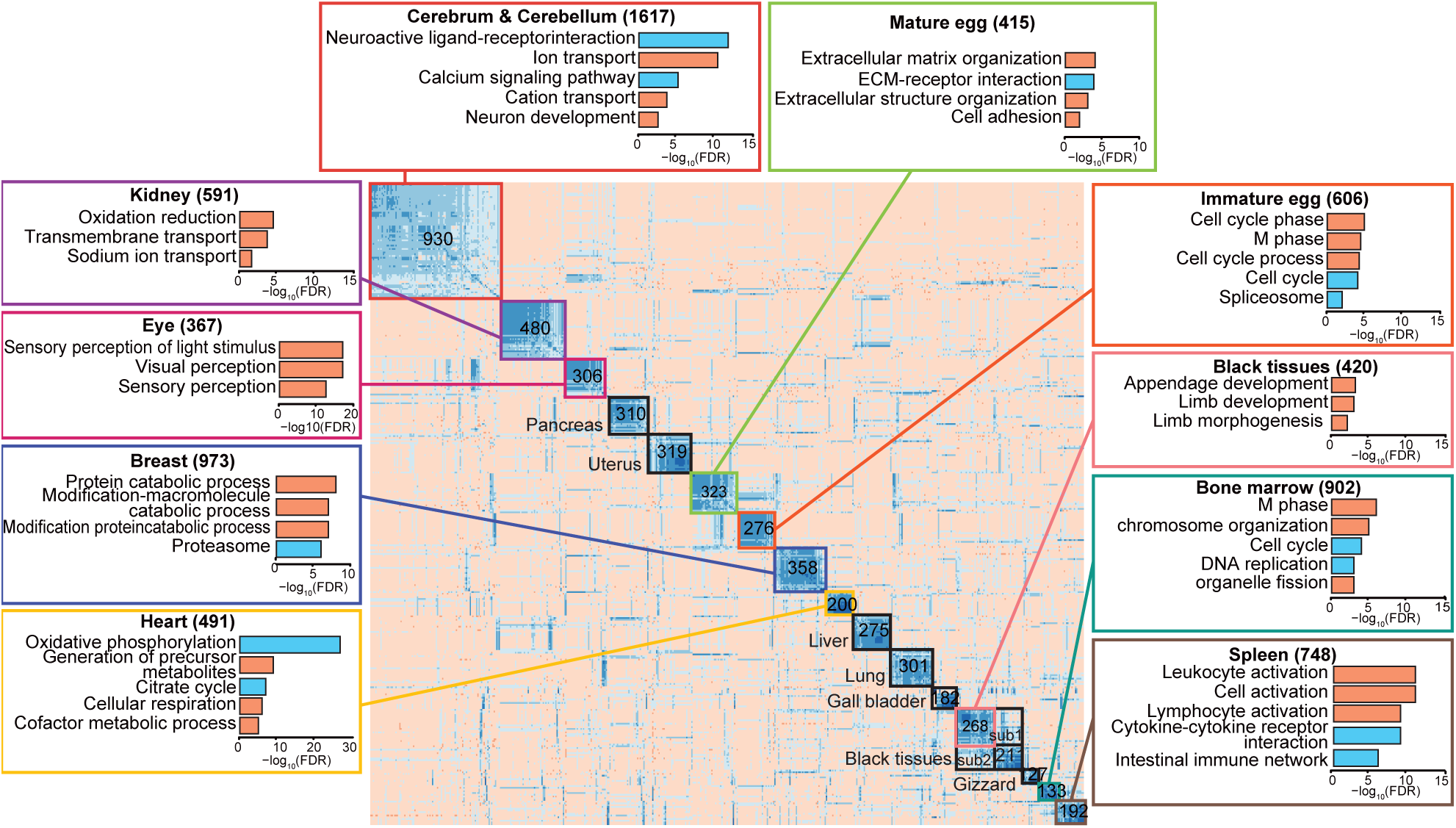
Co-expression clusters of lncRNAs and functional annotations. Co-expression clustering of lncRNAs across twenty tissues defines sixteen clusters and two sub-clusters specific to a tissue or a set of similar tissues. The boxes outlined in a color indicate clusters that have significant GO biological processes (orange bars) or KEGG pathway terms (cyan bars) associated with the protein-coding genes co-expressed with lncRNAs in the respective cluster. The significant enrichment of terms was tested using the hypergeometric test and adjusted by FDR, indicated with a logarithmic scale on the X-axis in the box. Clusters outlined in black are those that had neither a significant association with any GO term nor any co-expressed protein-coding genes. Sub-clusters in the clusters are indicated where appropriate. The number in each cluster indicates the number of lncRNAs in the cluster and the number in the boxes with functional terms indicates the number of co-expressed protein-coding genes.

The functional role of each co-expressed lncRNA cluster can be indirectly inferred by a set of co-expressed mRNAs ^48-50^. Thus, mRNAs that are exclusively co-expressed with each lncRNA cluster were identified with the following criteria: a mean Pearson’s correlation 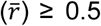 with members within a cluster and the differences between the corresponding 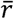 and the mean correlation 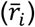 with all other groups ≥ 0.3, and were subsequently subjected to gene ontology (GO) analyses using DAVID ^60^ (Fig. 2; Supplementary table S3). In particular, 1617 mRNAs exclusively correlated to the brain-specific lncRNA group (930 lncRNAs) were identified and had been associated with brain-function specific terms, such as neuroactive ligand-receptor interaction (*q* = 2.18 X 10^−12^; False discovery rate, FDR correction). In contrast, 748 mRNAs exclusively correlated to spleen-specific lncRNAs were identified and had been associated with immune-related terms, such as leukocyte activation (*q* = 2.37X 10^−12^). Likewise, 10 out of 16 co-expression clusters of lncRNAs had functional evidence, with significantly enriched GO terms and KEGG pathways (Fig. 2).

### lncRNAs as epigenetic activators

The coherent expression of two different RNA classes could be in part the outcome of either active regulation by lncRNAs in *cis* and *trans*, or co-regulation by common regulators, such as TFs or epigenetic regulators, in *cis* and *trans* (Supplementary figure S3). Regulation of gene expression by lncRNAs often involves engagement with chromatin remodelers, such as polycomb repressive complexes (PRCs) that mediate the suppression of target mRNA expression ^61,62^ or demethylases that open the chromatin structure to enhance the expression of target mRNAs ^63,64^ (Supplementary figure S3a). Remote co-expression of lncRNAs and mRNAs can be also regulated by common TFs ^44,45^ (Supplementary figure S3b). Co-expressed genes tend to have common TF binding motifs in their promoters. However, *cis*-regulation of mRNA expression by lncRNAs is known to be associated with common epigenetic factors (Supplementary figure S3c) or enhancers (Supplementary figure S3d).

To find lncRNAs that act as epigenetic activators that reduce methylation levels, lncRNAs with expression levels that are significantly negatively correlated with the methylation level in the promoters of co-expressed protein-coding genes (nominal *P* ≤ 0.01) were examined in each co-expression cluster. In this case, the lncRNAs are thought to reduce the methylation level in the promoters of the co-expressed protein-coding genes. Of the lncRNAs in clusters, the expression of 15.0%~72.9% displayed significantly negative correlation with methylation levels in the promoters of co-expressed protein-coding genes, which were compared to those of random protein-coding gene cohorts (Fig. 3a). Clusters specific to brain, kidney, mature egg, breast, heart and spleen included significantly more lncRNAs with a significant correlation than did the random controls (*P* = 0.026~ 7.71 X 10^−13^) but this was not true for the black tissue cluster. To identify DNA methylation activators with more confidence, we also examined whether the expression and methylation of the co-expressed coding genes were correlated (nominal *P* ≤ 0.01). 820 lncRNAs in the clusters were identified as confident DNA methylation activator candidates (Fig. 3b). Genes encoding lncRNAs that act as DNA methylation regulators of protein-coding genes were mostly 100kb apart, and only five were within 100kb from target genes, suggesting that lncRNAs that function as epigenetic activators mostly play their roles in *trans*-form rather than *cis*-form.

**Figure 3.**
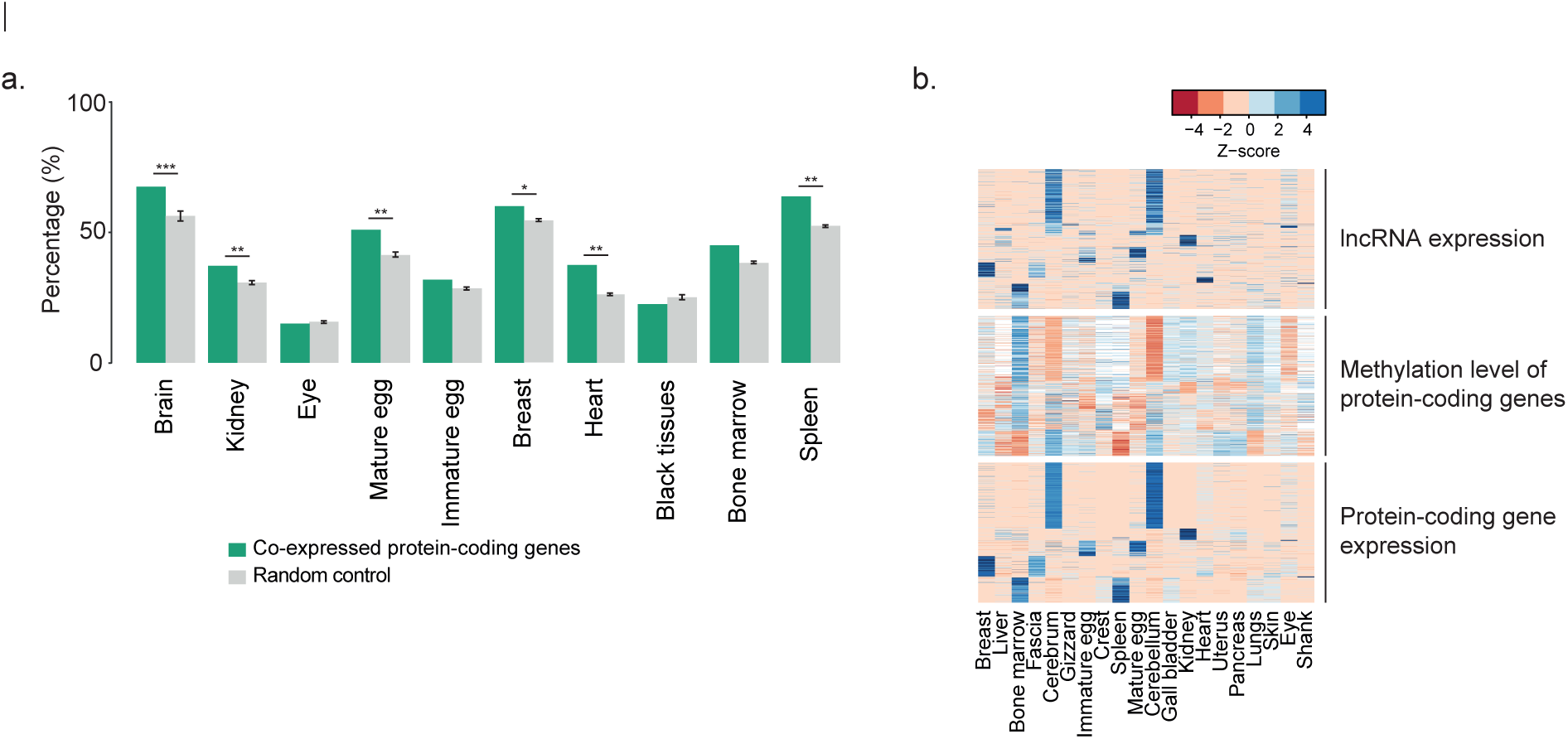
lncRNAs as epigenetic activators. **a**. The proportions of lncRNAs with expression levels that are correlated with the methylation level in the promoter of co-expressed protein-coding genes (dark green) in each cluster are shown in bar graphs. The numbers were compared to the mean methylation level of randomly selected protein-coding genes. To test the significance of the enrichment of lncRNAs as epigenetic activator candidates, 1000 number-matched random cohorts were compared to the original numbers (* *P* ≤ 0.05, ** *P* ≤ 0.01, *** *P* ≤ 0.001). **b**. lncRNAs as epigenetic activators whose expression levels are negatively correlated with the methylation level in the promoters of protein-coding genes, which in turn are negatively correlated with the level of protein-coding gene expression, as shown in heatmaps. The key indicates the z-score range of the expression values. White indicates N.A.

### Transcriptional regulation by common TFs

To identify co-expressed pairs of lncRNAs and mRNAs regulated by common TFs, TF binding sites (TFBSs) enriched in the promoters of the co-expressed genes were examined. For this analysis, sequences 2kb upstream of the co-expressed genes were extracted and enriched sequence motifs were identified using the multiple expectation-maximization for motif elicitation (MEME) suite ^65^ (see the “Prediction of TFBSs” section in the Methods for more details). The resulting motifs were subjected to analysis by the TOMTOM program ^66^ to annotate TFBSs based on TRANSFAC database v3.2 ^67^. As a result, 14 common TFs that have significantly abundant binding sites in the promoters of lncRNA and protein-coding genes were detected (Supplementary figure S4; corresponding to model 2). To discern TFs available in chicken genomes, PANTHER ^68,69^ was used to examine whether there are chicken orthologs of the TFs and whether the orthologs are expressed in the corresponding tissues (FPKM ≥ 1). Finally, five TFs, including HSF2 and SP1, were identified as candidates (Fig. 4a). HSF2 and SP1 binding sites were more recurrently detected across tissues than others and were significantly enriched in the promoters of 478 lncRNAs and 634 protein-coding genes. Although the binding motifs were slightly degenerated from the annotated motifs, the HSF2 motifs were similar in the promoters of lncRNA genes and protein-coding genes (Figs. 4b).

**Figure 4.**
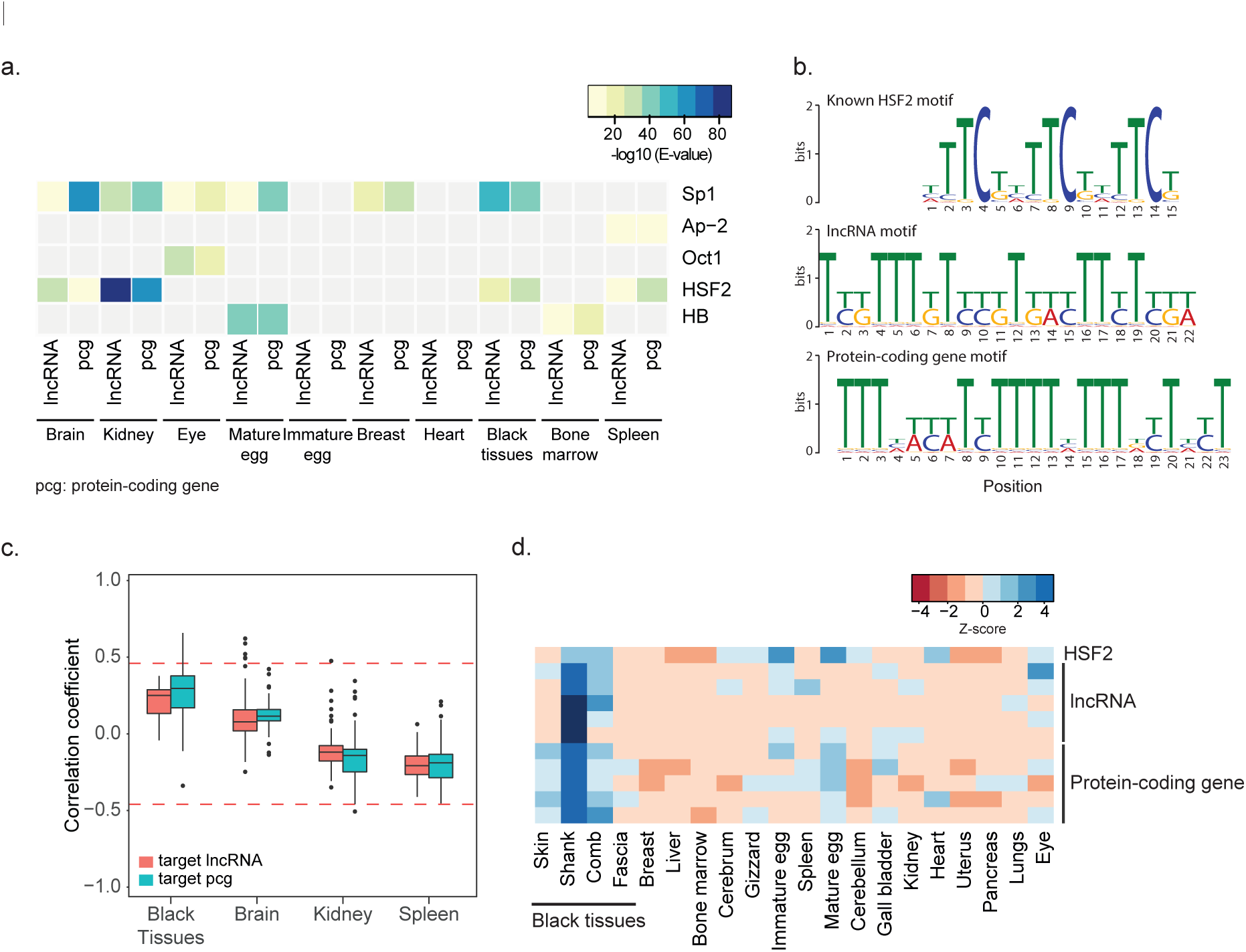
Co-transcriptional regulation of lncRNA and protein-coding genes by common TFs. **a**. TFs (Sp1, Ap-2, Oct1, HSF2, and HB) with binding motifs that are significantly co-enriched in the promoters of lncRNAs in a tissue-specific cluster and their co-expressed protein-coding genes are shown in the heatmap. The TFs are expressed in the indicated tissues. The significance of the motif enrichment was tested using MEME and E values are presented with color codes (blue: more significance, yellow: less significance) in the key. PCG indicates protein-coding gene. **b**. The HSF2 binding motif. A known motif is shown in the top panel, a motif in lncRNA promoters is shown in the middle panel, and a motif in protein-coding gene promoters is shown in the bottom panel. **c**. The expression correlation between co-regulated genes (red boxes for lncRNAs and green boxes for protein-coding genes) and HSF2 across tissues. Red lines indicate the significance level of the correlation coefficient (*P* ≤ 0.05). **d**. Expression pattern of HSF2 and its target genes that have the top 5 correlations with HSF2.

To examine further whether the respective TFs actually affect the expression of lncRNAs and protein-coding genes, the correlation between the expression of each TF and co-expressed genes in each cluster was examined. Interestingly, HSF2 expression had a strong positive correlation with expression of genes in black tissues but not in other tissues (Fig. 4c). The expression pattern for each of the five lncRNAs and protein-coding genes that were highly correlated with that of HSF2 was specific for skin, shank, and comb compared to other tissues (Fig. 4d). Thus, HSF2 is a promising candidate for regulating the black tissue-specific expression of lncRNAs and protein coding genes. Taken together, our data indicate that of a total of 3466 lncRNA in ten clusters, 615 (17.74%) appear to be co-regulated with co-expressed protein-coding genes by common TFs, such as HSF2.

### Coherent expression of neighboring lncRNA and protein-coding genes

Previous studies showed that lncRNAs and their neighboring protein-coding genes are highly correlated in their expression across tissues and developmental stages ^35,38^. To examine how the co-expressed lncRNAs and mRNAs in our study are co-localized in chromosomes, lncRNAs from each group were first classified based on the closest distances (≤10kb, ≤100kb, >100kb, and other chromosomes) from the significantly co-expressed protein-coding genes (nominal *P* ≤ 0.01; Pearson’s correlation) (Fig. 5a). Genes encoding co-expressed pairs of lncRNAs and mRNAs are significantly proximally co-localized within 10kb (Fig. 5a left; *P* ≤ 0.05, Fisher’s exact test), compared to random controls (Fig. 5a right) but not those of lncRNAs and mRNAs in the range of 10~100kb or in the 100kb outside. Overall, 2 ~ 15 % of the co-expressed pairs in the clusters tended to be proximally co-regulated within 10kb.

**Figure 5.**
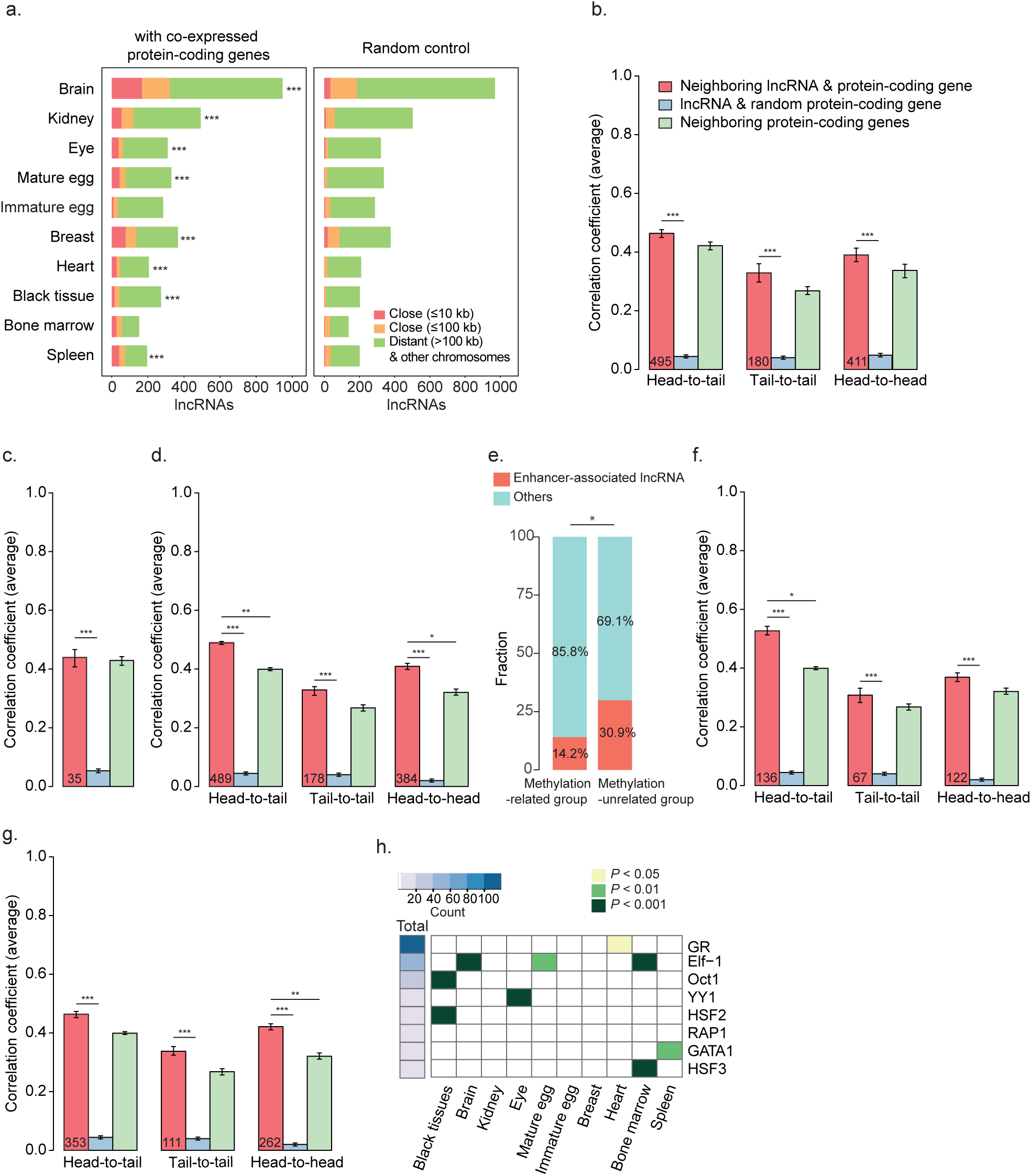
Co-regulation of neighboring lncRNA and protein-coding genes. **a**. Shown are the numbers of lncRNAs, classified by the distance from the closest protein-coding gene (red for the ≤ 10kb group, orange for the ≤ 100kb group, and green for the > 100kb or on another chromosome group) (left). *, **, and *** indicate *P* ≤ 0.05, ≤ 0.01, and ≤ 0.001, respectively. **b**. The average correlation coefficients of tissue-specific lncRNA and protein-coding gene pairs in close neighborhoods (≤ 10kb) are shown based on their relative orientations (head-to-tail, tail-to-tail, and head-to-head) (red bars). The average correlation coefficients of random pairs are also shown (blue bars) and those of tissue-specific protein-coding gene pairs in close neighborhoods (≤ 10kb) are shown with green bars. *, **, and *** indicate *P* ≤ 0.05, ≤ 0.01, and ≤ 0.001, respectively. Error bars indicate the standard error. The number in the bars indicates the number of analyzed pairs. **c**. The average correlation coefficients of neighboring lncRNA and protein-coding genes with similar methylation levels in their promoters (methylation-related) are shown in bar graphs. Otherwise, as in **b. d**. The average correlation coefficients of tissue-specific lncRNA and protein-coding genes (methylation-unrelated), except for those of **c**. Otherwise, as in **b. e**. The proportion of eRNAs (red) in the methylation-related group (**c**) and-unrelated group (**d**). ** indicates *P* ≤ 0.01. **f**. The average correlation coefficients of tissue-specific eRNAs. Otherwise, as in **b. g**. The average correlation coefficients of tissue-specific lncRNAs not associated with enhancers. Otherwise, as in **b. h**. TF binding motifs significantly associated with the eRNAs. The total count of the indicated TF binding sites in eRNAs is indicated in the heatmap (left) and the significance of the association over the total background is indicated with color-coded *P* values across tissues. The significance of a specific TF binding motif was tested using a binomial test in each tissue.

To examine how neighboring lncRNAs and protein-coding genes are tissue-specifically co-regulated, the pairs within 10kb were classified into three categories on the basis of their relative orientations (head-to-tail, tail-to-tail, or head-to-head). The correlation coefficients of the pairs in each category were compared to those of lncRNA and random protein-coding gene controls from tissue-specific gene sets (Fig. 5b) or from ubiquitously expressed gene sets (Supplementary figure S5a). Both neighboring lncRNA and protein-coding gene pairs displayed significantly greater correlation than did random controls, regardless of the category, in both sets (Fig. 5b; Supplementary figure S5a). The correlations were also compared to those of neighboring protein-coding gene pairs. Whereas the correlations of the ubiquitously expressed, neighboring lncRNAs and protein-coding genes were significantly lower than those of ubiquitously expressed neighboring protein-coding gene pairs in the head-to-tail and head-to-head categories (Supplementary figure S5a), the correlation coefficients of the tissue-specific pairs were slightly yet insignificantly higher than those of neighboring protein-coding gene pairs (Fig. 5b).

To dissect factors that affect the co-regulation of tissue-specific neighboring lncRNA and protein-coding gene pairs, the pairs with a high correlation (*P* ≤ 0.05) between the methylation levels of their promoters (methylation-related group – model 3) and those with no correlation (methylation-unrelated group) were divided. Tissue-specific neighboring lncRNA and protein-coding gene pairs showed no more expression correlation than did neighboring protein-coding genes in the methylation-related group (Fig. 5c; *P* = 0.71, Wilcoxon rank sum test), whereas they did show a significantly higher correlation in the methylation-unrelated group (Fig. 5d; *P* ≤ 0.001 for head-to-tail, *P* ≤ 0.05 for head-to-head, Wilcoxon rank sum test), which suggests that neighboring lncRNAs and protein-coding genes in the methylation-unrelated group have a regulatory interaction between them.

### Enhancer-associated RNA-mediated gene regulation

Previous studies showed that lncRNAs associated with enhancers could regulate their neighboring protein-coding genes ^70^. Genomic association between lncRNAs and enhancers, detected in embryonic developmental stages in the chicken ^71^, revealed that lncRNAs in the methylation-unrelated group are more significantly associated with enhancers than those in the other group (Fig. 5e; *P* = 2.72 X 10^−6^; Fisher’s exact test). As a result, 136 head-to-tail lncRNAs, 67 tail-to-tail lncRNAs and 124 head-to-head lncRNAs were considered as enhancer-associated lncRNA candidates (eRNAs; Supplementary Table S4). The eRNAs (corresponding to model 4) had a greater correlation with neighboring protein-coding genes only in the head-to-tail group (Fig. 5f), whereas non-eRNAs displayed a greater correlation in the head-to-head orientation, which could allow sharing of common promoters (Fig. 5g). A few eRNAs were discovered to have strong bi-directional transcriptional activity (Supplementary figure S5b; see the “Transcriptional activity of eRNAs” section in the Methods for more details), as previously reported ^72,73^

Next, to identify TFs binding to genomic regions that transcribe eRNAs, TF binding sites detected from all the genomic regions associated with enhancers were profiled and were compared to those of TFs detected from the enhancers specific to a certain tissue (Fig. 5h). Oct1 and HSF2 binding sites were significantly localized in eRNAs specific to black tissues (*P* < 3.09 X 10^−5^ for Oct1; *P* < 3.11 X 10^−7^ for HSF2; binomial test). Besides the TFs specific to black tissues, GR, YY1, RAP1 and GATA1, and HSF3 binding sites were localized in eRNAs specific to heart, eye, spleen and bone marrow, respectively (Fig. 5h). Interestingly, HSF2 was a common TF candidate for co-regulating lncRNAs and protein-coding genes at a distance (Fig. 5d).

### Conserved black skin-specific lncRNAs

As mentioned earlier, unlike other chicken breeds, both the plumage and skin of the Ogye are black. To identify lncRNAs potentially functionally related to this trait, lncRNAs specifically co-expressed in black tissues (Fig. 2) were further investigated by comparing to those in non-black skin of other chicken breeds. Of 479 lncRNAs specific to black tissues, 47 were significantly two-fold up-(29) or down-regulated (18) in Ogye black skin, compared to those in brown leghorn skin (Fig. 6a; Supplementary table S5; FDR < 0.05).

**Figure 6.**
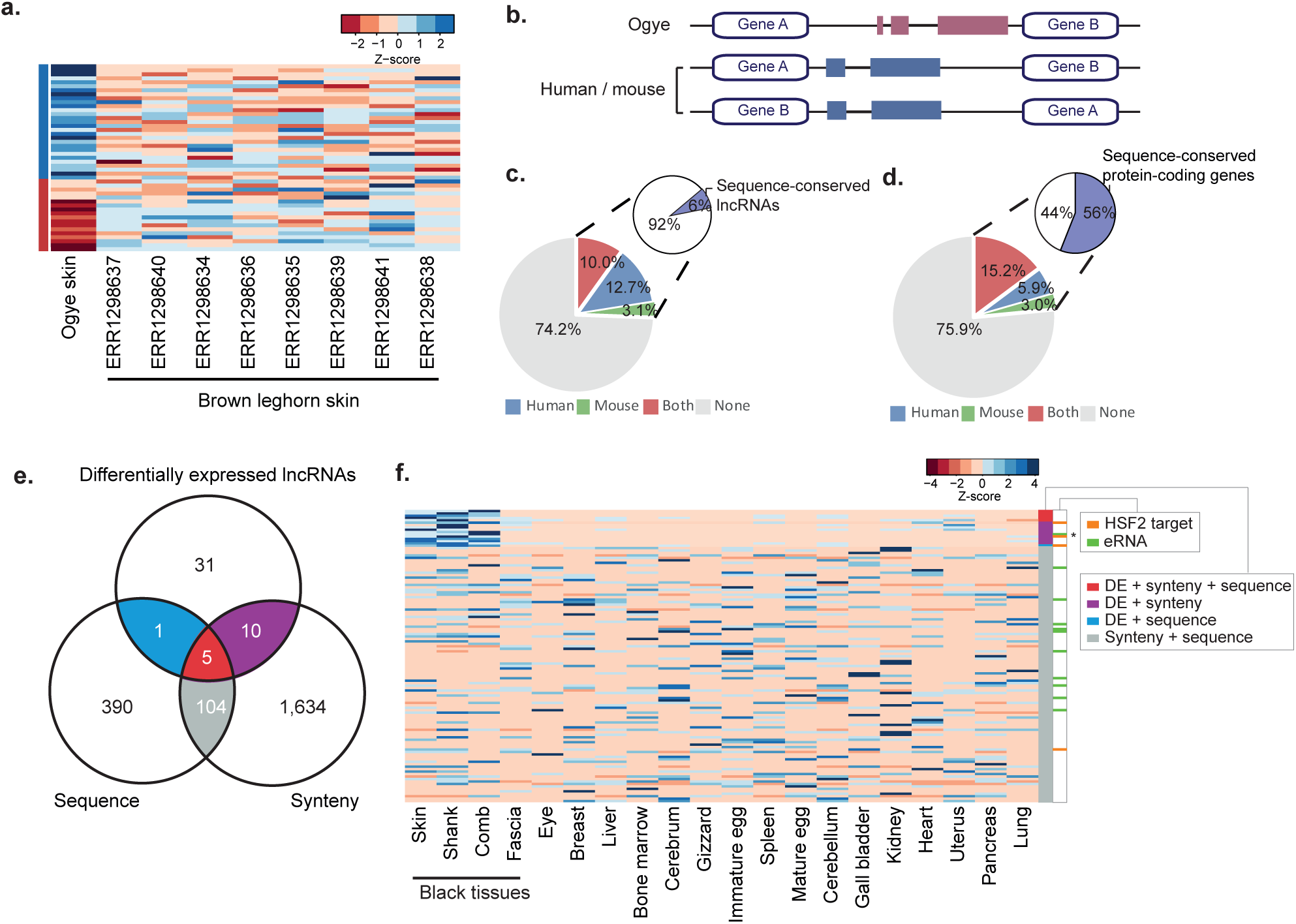
Black tissue-specific lncRNAs with sequence and synteny conservation. **a**. The expression patterns of differentially expressed lncRNAs in Ogye skin, compared to brown leghorn skin samples. Expression levels are indicated with a color-coded Z-score (red for low and blue for high expression) as shown in the key. **b**. A cartoon showing a lncRNA that is syntenically conserved with up-and down-stream protein-coding genes in the human and/or mouse genome. **c**. The fraction of lncRNAs with syntenic conservation in the human (blue), mouse (green) or both (red) genomes is shown in the pie chart. Of the syntenically conserved lncRNAs, the fraction of lncRNAs with sequence conservation (purple) in the human or mouse genome is indicated in the secondary pie charts. **d**. The fraction of protein-coding genes with synteny conservation is indicated in the pie chart. Otherwise, as in **c. e**. The numbers of differentially expressed lncRNAs in black skin with evidence of sequence and synteny conservation are indicated in a Venn diagram. **f**. Evidence for differential expression (DE) + synteny + sequence (red), DE + synteny conservation (purple), or DE + sequence conservation (blue) for 16 black-skin specific lncRNAs is shown in a heatmap. 104 non-specific lncRNAs with evidence of sequence + synteny conservation are indicated in gray. The co-regulation models associated with a certain lncRNA are indicated to the left with color codes (orange for HSF2 binding and green for eRNAs). * indicates the eRNA associated with HSF2. The expression level is indicated with a color-coded z-score, as shown in the key.

To find functionally conserved lncRNAs, the 47 differentially expressed lncRNAs were examined for synteny and sequence conservation in human and mouse genomes. Synteny conservation considers whether orthologs of a certain lncRNA’s neighboring genes are positionally conserved in these mammalian genomes (Fig. 6b). As a result of this analysis, about 10% of lncRNAs were found to be syntenically conserved in both the human and mouse genomes and about 25% were syntenically conserved in at least one genome (Fig. 6c; Supplementary table S6), percentages that are comparable to those of the protein-coding genes (Fig. 6d). However, sequence similarity analyses by the BLAST showed that only 6% of the syntenically conserved lncRNAs had conserved sequences relative to sequences in either the human or mouse genomes (Fig. 6c; Supplementary table S6), which is much lower than that of protein-coding genes (56%). Taken together, our data showed that 16 lncRNAs were syntenically or sequentially conserved and differentially expressed in black tissue (Fig. 6e).

Of the 16 lncRNAs that have evidence of black tissue-specific function, four, including eRNAs, were associated with HSF2 binding motifs, whereas of the 104 that have synteny and sequence conservation but are not differentially expressed in black tissues, only one was associated with HSF2. The presence of HSF2 binding motifs appears to be significantly related to black tissue-specific expression (Fig 6f; *P* ≤ 0.0008, Fisher’s exact test). For instance, linc-THEM184c is significantly up-regulated in black tissue (Fig. 7b), its locus is syntenically conserved with neighboring genes, TMEM184C and EDNRA, in both human and mouse genomes, and its promoter includes a HSF2 binding motif (Fig. 7a). In addition, the protein-coding genes that are co-expressed with this lncRNA are enriched for GO terms that are functionally relevant for black skin: keratinocyte differentiation, angiogenesis, and ECM-receptor-interaction (Fig. 7c). Among the co-expressed genes, 31 have HSF2 binding sites in their promoters (Fig. 7a). As another example, black-tissue specific linc-FAM204A is syntenically conserved with the RAB11FIP2 and FAM204A genes in the human and mouse genomes (Supplementary figure S6a). This lncRNA was highly expressed in black tissues including the skin, shank, and comb but had no expression in other tissues except for the eye (Supplementary figure S6b). The co-expressed protein-coding genes are enriched for functionally relevant GO terms melanogenesis, ECM-receptor interaction, and Wnt signaling (Supplementary figure S6c). Interestingly, the human and Ogye lncRNA orthologs share a conserved sequence of 389 nt, which includes multiple miRNA 7-mer target sites (Supplementary figure S6a).

**Figure 7.**
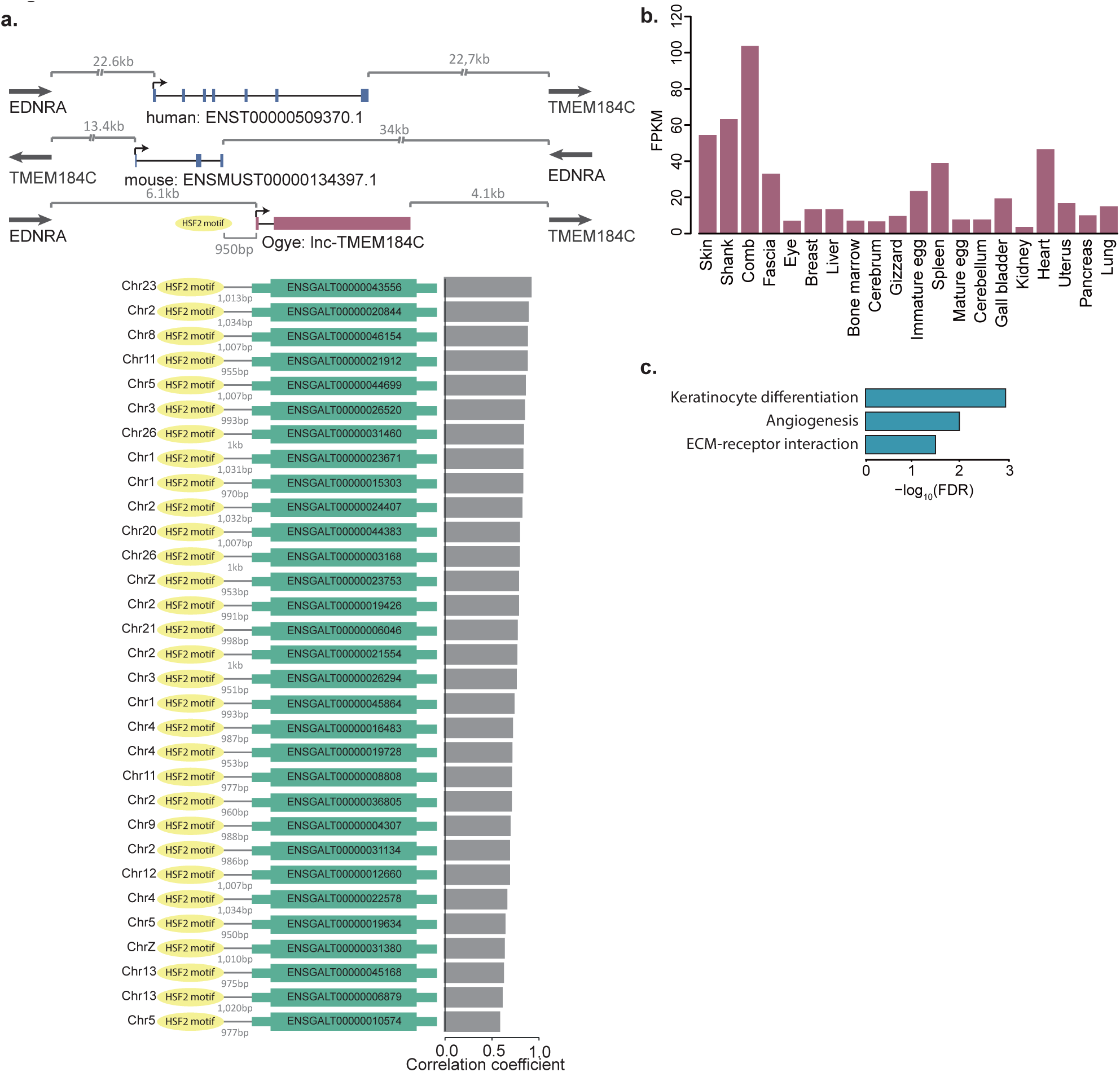
An example of black skin-specific lncRNAs with synteny conservation, which is transcriptionally regulated by HSF2. **a**. Ogye lncRNA (lnc-TMEM184C) with synteny conservation in human and mouse genomes (top). The lncRNA has an HSF2 binding motif in its promoter; this motif is also present in the promoters of protein-coding genes with correlated expression (below). Gray bar plots indicate the expression correlation between the lncRNA and the protein-coding genes. **b**. The lnc-TMEM184C expression pattern across 20 tissues. **c**. GO terms that are significantly associated with the protein-coding genes that are co-expressed with lnc-TMEM184C.

## Discussion

The majority of lncRNAs showed a tissue-specific expression pattern, defining functionally coherent co-expression clusters. The tissue-specific expression and the coherent expression of lncRNA genes with other protein-coding genes could be attributed to common epigenetic and transcriptional regulation. In fact, of the lncRNAs in clusters, 39.3% had evidence associating them with at least one model (Supplementary figure S7a); most commonly, these involved lncRNAs that act as epigenetic activators of protein-coding gene expression and common TFs that bind to the lncRNA and protein-coding gene promoters (Supplementary figure S7b). Interestingly, 126 lncRNAs had evidence supporting both the epigenetic activator and TF models (Supplementary figure S7c). 79 lncRNAs had functional evidence supporting their identity as eRNAs. Although lncRNAs are known to be mostly involved in epigenetic repression of genes, our study intentionally focused on lncRNAs as epigenetic activators by correlating the level of lncRNAs and the methylation in target gene promoters. Furthermore, because only a subset of CpG sites are sometimes related to the chromatin state and transcriptional activity of target genes, averaging CpG methylation signals in the promoter might underestimate the fraction of epigenetically activating lncRNAs in our study.

Although protein-coding genes co-expressed with lncRNAs in black tissues seem to be not associated with epigenetic regulation by DNA methylation (Fig. 3a), lncRNA and protein-coding genes co-expressed in black tissues had HSF2 binding sites in their promoters and were specifically correlated with the level of HSF2 across tissues, supporting that the genes are co-regulated by HSF2 (Fig. 4). Moreover, enhancers that included HSF2 binding sites were associated with eRNAs specific to black tissue (Fig. 5h), indicating that HSF2 is the most likely regulator of black tissue-specific expression. Because the ancestor of Ogye appears to have originated in the rainforest, it makes sense that heat shock-related factors could be involved in melanogenesis and hyper-pigmentation processes, which would help avoid the absorption of too much heat. One of the black skin-specific lncRNAs, lnc-THMEM184c, is most abundantly expressed in comb, and HSF2 appears to co-regulate lnc-THMEM184c and its co-expressed protein-coding genes, which are related to keratinocyte differentiation and ECM-receptor interaction (Fig. 7).

In addition, several previous studies that also focused on animal coat color showed that the color can be determined by the amount and type of melanin produced and released by melanocytes present in the skin ^74,75^. Melanin is produced by melanosomes, large organelles in melanocytes, in a process called melanogenesis. Wnt signaling has a regulatory role in the melanogenesis pathway and is also required for the developmental process that leads to melanocyte differentiation from neural crest cells ^76,77^. One of the candidate lncRNAs related to the process is linc-FAM204A, whose co-expressed protein-coding genes are associated with GO terms melanogenesis, ECM-receptor interaction, and Wnt signaling pathway (Supplementary figure S6c). linc-FAM204A, which contains a short-conserved motif, is broadly preserved in mammalian genomes, including the human, rhesus macaque, mouse, dog, and elephant genomes. Among these orthologs, the human ortholog is known as CASC2, and is suppressed in lung, colorectal, renal and other cancers by miR-21-5p targeting via the conserved 7-mer site (Supplementary figure S6a).

Taken together, these results indicate that coding and non-coding RNAs functionally relevant to black and other tissues could help explain unique genomic and functional characteristics of a Korean domestic chicken breed, Yeonsan Ogye. Additionally, these findings could provide unprecedented insight for future studies with industrial and agricultural applications, as well as for scientific analysis of chicken genomes.

## Methods

### Acquisition and care of Yeonsan Ogye

Yeonsan Ogye chickens (object number: 02127), obtained from the Animal Genetic Resource Research Center of the National Institute of Animal Science (Namwon, Korea), were used in the study. The care and experimental use of Ogye was reviewed and approved by the Institutional Animal Care and Use Committee of the National Institute of Animal Science (IACUC No.: 2014-080). Ogye management, treatment, and sample collection and further analysis of all raw data were performed at the National Institute of Animal Science.

### Datasets

To profile expression of protein-coding genes in the Ogye genome, *Gallus gallus* (red junglefowl) protein-coding genes were downloaded from Ensembl biomart (release 81; http://www.ensembl.org/biomart), and were mapped onto the Ogye draft genome v1.0 using GMAP (v2015-07-23)^78^. Genes that had greater than 90% coverage and identity were selected as Ogye protein-coding genes. As a result, 14,264 protein-coding genes were subjected to further analysis.

Total RNA samples and Bisulphite-treated DNA samples were collected from twenty different tissues (Breast, Liver, Bone marrow, Fascia, Cerebrum, Gizzard, Immature egg, Comb, Spleen, Mature egg, Cerebellum, Gall bladder, Kidney, Heart, Uterus, Pancreas, Lung, Skin, Eye, and Shank) from 8-month-old Ogye (Supplementary figure S1a). About 1.5 billion RNA-seq reads (843 million single-end reads and 638 million paired-end reads) and 123 million RRBS reads were analyzed (Fig. 1a).

### Tissue-specific, differentially methylated CpG sites

RRBS reads were aligned to the Ogye draft genome (v1.0)^56^ using Bismark ^79^. The methylation level of each cytosine in a CpG region was calculated using Bismark methylation extractor. A tissue-specific, differentially methylated CpG (tDMC) site is defined as one in which its mean methylation across tissues is at least four time greater than the minimum signal in a certain tissue. The tDMC sites were found in the promoter region of each gene, defined as the region 2 kb upstream of the 5’ end of genes.

### Expression profiling

The expression values (FPKM) of lncRNA and protein-coding genes were estimated using RSEM (v1.2.25) in each tissue. The values across tissues were normalized using the quantile normalization method. In all downstream analyses, lncRNA or protein-coding genes with FPKM ≥ 1 in at least one tissue were used. lncRNAs for which the maximum expression value across twenty tissues was at least four-fold higher than the mean value were considered to exhibit tissue-specific expression.

### Phylogenetic analysis of expressed lncRNAs across tissues

To perform phylogenetic analysis of commonly expressed lncRNA genes among tissues, the list of expressed lncRNAs in each tissue was used as a input vector for phylogenetic clustering. The clustering was done using the PHYLIP package. lncRNAs with FPKM ≥ 1 in a certain tissue were considered to be expressed in a certain tissue. As two tissues share more common genes, they become more closely clustered.

### Clustering of co-expressed lncRNAs

Hierarchical clustering was performed to search for expression clusters of lncRNAs across twenty tissues using Pearson’s correlation coefficient metrics. Clusters in which more than 80% of their members are most highly expressed in the same or related tissues (brain and black tissues) were regarded as tissue-specific. Sub-clusters in the brain and black tissue clusters were further defined with the same criterion mentioned above.

### Defining coding genes co-expressed with lncRNAs in a cluster

Protein-coding genes with a high mean correlation with lncRNAs in a cluster (Pearson’s correlation ≥ 0.5), but for which the mean correlation to the cluster is at least 0.3 greater than those of other clusters, were assigned to the co-expressed set of the cluster. Each set of mRNAs was used to perform gene ontology (GO) term and pathway enrichment analyses using DAVID ^60^. Terms were only selected when the false discovery rate (FDR) *q* value was ≤ 0.05.

### Correlation of the methylation level of neighboring lncRNA and protein-coding genes

The methylation levels at CpG sites in the promoters of neighboring lncRNA and protein-coding genes were correlated with each other over twenty tissues (using Pearson’s correlation coefficients). Only tissues in which a certain position had sufficient read coverage (at least five) were considered for measuring the correlation. If the nominal *P* value was ≤ 0.05, then the pair of lncRNA and protein-coding genes was considered as having a significantly correlated interaction.

### Correlating the expression level of lncRNAs with the methylation level of protein-coding genes

To identify lncRNAs as potential epigenetic activators, the expression of lncRNAs and the methylation at CpG sites in the promoters of protein-coding genes were correlated over twenty tissues using a non-parametric correlation method (Spearman’s correlation). Only pairs of lncRNA and protein-coding genes exhibiting a nominal *P* value ≤ 0.01 were considered as having a significantly correlated interaction. Of the resulting pairs, if the protein-coding mRNAs had a significant correlation (nominal *P* value ≤ 0.01) between their expression level and the methylation level in their promoter, its paired lncRNA was regarded as an epigenetic activator.

### Prediction of TFBSs

To identify enriched TFBSs in the promoters of the co-expressed lncRNAs in each tissue-specific cluster and in the promoters of the co-expressed protein-coding genes within the cluster, the promoter sequences were examined using the MEME suite (V4.9.0). Motifs that exhibit an E-value ≤ 1 X 10^−5^ were selected as enriched motifs, associated with the corresponding tissue. The resulting motifs were searched for in the Transfac database ^67^ using TomTom ^66^.

### Identification of enhancer regions

To annotate enhancer regions in the Ogye draft genome, annotation files including all enhancers in the *Gallus gallus* (*red junglefowl*) genome were downloaded from the NCBI gene expression omnibus (GEO, GSE75480). Enhancer sequences extracted using our in-house script were aligned to the Ogye draft genome using BLAST (-p blastn). Regions that significantly matched the original enhancers (E-value ≤ 1 X 10^−5^) and with high coverage of more than 80% were annotated as Ogye enhancers.

### Transcriptional activity of eRNAs

To examine bi-directional transcriptional activity of eRNAs, total mapped reads in the range spanning 1kb upstream to 1 kb downstream of the eRNA transcription start site (TSS) were re-examined on both forward and reverse strands.

### Correlation of expression between neighboring lncRNA and protein-coding genes

Pairs consisting of a lncRNA and its closest neighboring protein-coding gene within 10kb were classified into three groups based on their genomic orientations: head-to-head (can be divergently overlapped), head-to-tail (including only independent lncRNAs with evidence of a TSS and cleavage and polyadenylation site; otherwise, these lncRNAs must be at least 1kb apart from each other), and tail-to-tail (can be convergently overlapped). The correlation of the expression of these pairs was calculated over twenty tissues using Pearson’s correlation method. The average correlation coefficient values and their standard errors were calculated in the respective groups. As a random control, the expression of 1000 random pairs of lncRNA and protein-coding genes were correlated using the same method. As another control, number-matched pairs of neighboring protein-coding genes were also correlated with each other.

### Synteny and sequence conservation

To examine the conservation of synteny of a lncRNA, its closest downstream and upstream neighboring protein-coding genes in the Ogye genome were matched to their orthologous genes in the mouse and human genomes. If a lncRNA is located between the two orthologous genes, regardless of direction, that lncRNA was regarded as syntenically conserved. GENCODE lncRNA annotations (v25 for human and vM11 for mouse) were analyzed for this study. To check for sequence conservation, Ogye lncRNA sequences were aligned to lncRNA sequences from other species, intronic sequences, and their flanking sequences (up to 4 Mb) using BLAST. For a significant match, an E-value 1 X 10^−6^ was used as a cutoff.

### Analysis of lncRNA differential expression

To identify lncRNAs that are differentially expressed between Ogye and Brown leghorn skin tissues, Brown leghorn skin RNA-seq data were downloaded from the NCBI SRA (ERR1298635, ERR1298636, ERR1298637, ERR1298638, ERR1298639, ERR1298640, and ERR1298641). Reads were mapped to the *Gallus gallus* Galgal4 reference genome using Bowtie (V1.0.0), and the average mismatch rates were estimated across read positions. If the mismatch rate was greater than 0.1 at a certain position, sequences on high mismatch side of the position were trimmed using seqtk (https://github.com/lh3/seqtk), and then sickle was used with the default option for read quality control. Preprocessed reads from RNA-seq data were mapped onto the chicken Galgal4 reference genome using STAR. The read counts of lncRNAs were performed using HTSeq (v0.6.0) and the differential expression analysis was performed using DESeq ^80^. Genes with a greater than two-fold difference in expression and a FDR *q* value ≤ 0.05 were considered to be differentially expressed.

## Acknowledgements

We thank all members of the BIG lab for helpful comments and discussions.

## Author’s contributions

JWN and HHC conceived and supervised the project. CYC provided materials. DJL, KTL, and YJD performed RNA-seq experiments. HSH performed the analyses. JWN and HSH wrote the manuscript. All authors read and approved the final manuscript.

## Competing financial interests

The authors declare no competing financial interests.

## Data availability

Raw RNA-seq and RRBS data from twenty different Ogye tissues have been submitted to the NCBI Gene Expression Omnibus (GEO; https://www.ncbi.nlm.nih.gov/geo/; GSE104351) under SuperSeries accession number GSE104358. All lncRNA catalogs and expression tables from this study have also been submitted to NCBI GEO (GSE104351) under the same SuperSeries accession number, GSE104358.

## Funding

This work was supported by the Program for Agriculture Science & Technology Development (Project No. PJ01045301; PJ01045303) of the Rural Development Administration.

## Supplementary Information

Supplementary Figures S1-S7.

Supplementary Table S1. Tissue-specific lncRNAs and ubiquitously expressed lncRNAs.

Supplementary Table S2. lncRNAs corresponding to each cluster (16 clusters).

Supplementary Table S3. Enriched biological process GO terms and KEGG pathways for protein-coding genes co-expressed with each cluster (FDR ≤ 0.05).

Supplementary Table S4. lncRNAs associated with enhancers and neighboring protein-coding genes in terms of their genomic orientations: head-to-tail, tail-to-tail, head-to-head.

Supplementary Table S5. lncRNAs that are differentially expressed between Ogye and Brown leghorn skin.

Supplementary Table S6. Synteny-conserved lncRNAs and sequence-conserved lncRNAs.

